# Recent adaptation in an imperiled salmonid revealed by museum genomics

**DOI:** 10.1101/2024.04.25.590849

**Authors:** Andrew G. Sharo, Megan A. Supple, Randy Cabrera, William E. Seligmann, Samuel Sacco, Cassondra D. Columbus, Devon E. Pearse, Beth Shapiro, John Carlos Garza

**Affiliations:** Department of Ecology and Evolutionary Biology, University of California, Santa Cruz, CA, USA; Institute of Marine Sciences, University of California, Santa Cruz, CA, USA; Southwest Fisheries Science Center, National Marine Fisheries Service, National Oceanic and Atmospheric Administration, Santa Cruz, CA, USA; Howard Hughes Medical Institute, Santa Cruz, CA, USA; Department of Ocean Sciences, University of California, Santa Cruz, CA, USA

## Abstract

Steelhead/rainbow trout (*Oncorhynchus mykiss*) is an imperiled salmonid with two main life history strategies: migrate to the ocean or remain in freshwater. Domesticated hatchery forms of this species have been stocked into almost all California waterbodies, possibly resulting in introgression into natural populations and altered population structure.

We compared whole-genome sequence data from contemporary populations against a set of museum population samples of steelhead from the same locations that were collected prior to most hatchery stocking.

We observed minimal introgression and few steelhead-hatchery trout hybrids despite a century of extensive stocking. Our historical data show signals of introgression with a sister species and indications of an early hatchery facility. Finally, we found that migration-associated haplotypes have become less frequent over time, a likely adaptation to decreased opportunities for migration. Since contemporary migration-associated haplotype frequencies have been used to guide species management, we consider this to be a rare example of shifting baseline syndrome that has been validated with historical data.

We suggest cautious optimism that a century of hatchery stocking has had minimal impact on California steelhead population genetic structure, but we note that continued shifts in life history may lead to further declines in the ocean-going form of the species.

## Introduction

Steelhead/rainbow trout (*Oncorhynchus mykiss*) is a large cold-water fish that has evolved two distinct life histories. After juveniles develop in their natal streams, some individuals migrate to the ocean to mature, as is typical in many salmonid species. Those individuals that migrate are termed steelhead, and adult steelhead later return to freshwater to reproduce. This migratory life-history strategy is termed anadromy. In contrast, some individuals from the species remain in freshwater for their entire life cycle and then reproduce in their natal waterbody. These resident fish are termed rainbow trout.

In much of their native range in the rivers and streams of Western North America, steelhead are declining due to habitat loss and anthropogenic alterations to freshwater and marine ecosystems^1^. Despite these challenges, the species is commonly fished for recreation and is one of the most cultivated aquaculture species worldwide. Historically, native peoples relied upon abundant and dependable steelhead populations for sustenance. Yet today, many populations, particularly those in California, are protected by the US Endangered Species Act^2^.

As steelhead/rainbow trout populations began to decline at the end of the 19th century, the state of California’s Department of Fish and Wildlife (CDFW) sought to improve fishing opportunities. Through a state-wide system of hatcheries, CDFW has reared and released billions of rainbow trout in California since the early 1900s. Although some hatcheries released trout reared from eggs taken from the same waterbody, most eggs were produced from domesticated strains that were developed from distant populations, including stocking trout domesticated from inland populations^3^ into coastal California waterways.

When domesticated animals are released into natural populations of conspecifics, there are several possible outcomes: no effect^4^, introgression^5^, or even complete replacement^6^. Typically fitness decreases when native taxa are introgressed by invasives^7,8^. When a rare group is put in contact with a more common group, the rare group may lose its distinct ancestry and population dynamics until it succumbs to what is sometimes called extinction by hybridization^9,10^.

Despite previous research, the extent of hatchery introgression in protected California steelhead populations is unclear. Clemento et al. used neutral multi-allelic genetic marker (microsatellite) data and found no clear signs of introgression when fish above and below barriers were compared^11^. Similarly, Garza et al. examined population structure in steelhead from migratory populations in coastal California with the same set of microsatellite markers and found strong isolation by distance, i.e., geographic and genetic distance were correlated^12^. Conversely, Pearse et al. sequenced a short segment of mitochondrial DNA (mtDNA) from historical population samples of steelhead from museum collections taken in several California streams, including one that overlapped with Clemento et al.’s study area, and found reduced isolation by distance in modern populations compared with historical populations, which was attributed to hatchery rainbow trout introgression and population fragmentation^13^.

It is also unclear whether protected steelhead populations below barriers have experienced selection against anadromy due to their impeded migratory routes. Anadromy has been found to be strongly associated with a ∼55 Mb double inversion on chromosome Omy05 in California populations^14^. Previous studies suggest that above-barrier fish with the migration-associated haplotype are more likely to pass below barriers, generating purifying selection that results in a greater proportion of resident fish above barriers^15^, but effects on below-barrier fish are unknown.

This question of whether anadromy has been selected against in below-barrier steelhead populations may be an example of shifting baseline syndrome, first coined by Pauly^16^. Prior to human impact, there was a pre-disturbance baseline level of the migration-associated haplotypes in steelhead populations. However, this baseline is unknown, and so researchers have developed an estimated baseline from contemporary populations, and used this to propose potential restoration scenarios^17,18^. A recent study found that the migration-associated haplotype ranged from 25–96% frequency in below-barrier California populations^15^. However, whether this reflects the true, pre-disturbance baseline or a more recent state can only be answered by analyzing historical populations.

Recently, progress in DNA sequencing of museum specimens^19,20^ have made it possible to extract meaningful genomic data from formaldehyde-preserved specimens^21^, including museum samples of fish. We apply these advances to sequence whole nuclear genomes (>1X coverage for 92% of historical samples; Fig. S1) from a rare set of population samples taken from California steelhead around the turn of the 20th century (prior to most hatchery stocking), and archived in the National Museum of Natural History. We compare these data with contemporary populations from five rivers in northern and central California (Fig. 1A-C, Table S1), along with five hatchery strains, to assess temporal changes in population structure, hatchery introgression, and migration-associated loci.

**Fig. 1.**
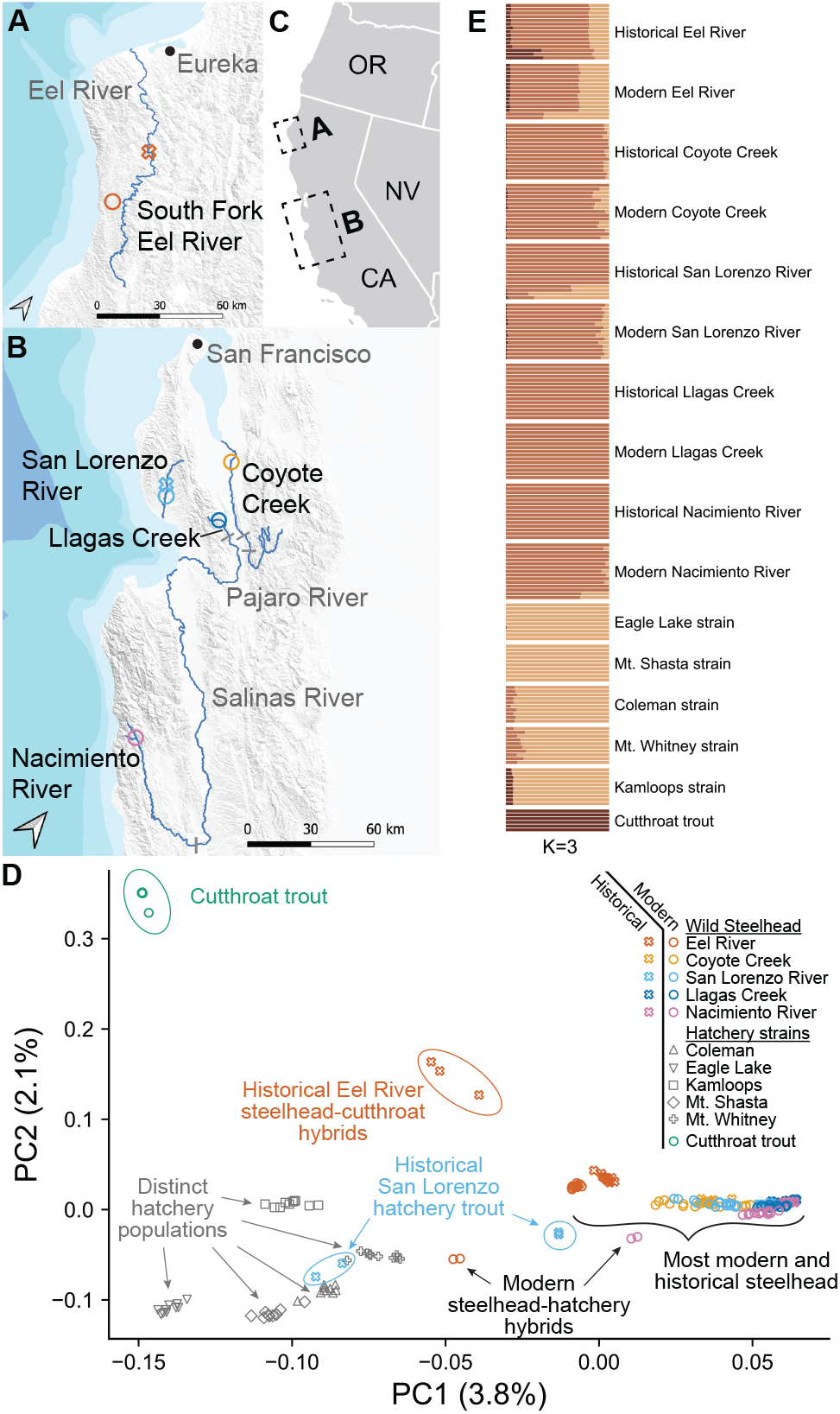
Sampling scheme and population genetic analysis. (**A-C**) Locations where fish were sampled from California rivers. Cities and mainstems of sampled tributaries are written in gray. Each marker represents a sampling location, where X’s are historical and O’s are contemporary, colored according to the legend in (D). For some rivers, exact historical sampling location is unknown. Gray bars indicate impassable barriers. (**D**) PCA analysis of genomic data with distinct colors and symbols indicating separate geographic populations and time periods. Hatchery populations are in gray. (**E**) Unsupervised ADMIXTURE (K = 3 clusters) analysis. Each bar represents an individual and each color represents a different ancestry assignment.

## Results

### Contemporary steelhead derive most of their ancestry from historical steelhead rather than hatchery strains or cutthroat trout

The broad population structure of California steelhead has been largely unaffected by billions of stocked hatchery trout. We carried out principal component analysis (PCA) on the historical, contemporary, and hatchery steelhead/rainbow trout genomes, together with previously generated WGS data from a sister species, cutthroat trout (*O. clarkii*). To avoid bias due to coverage differences and damage in historical DNA, we called pseudo-haplotypes and analyzed only transversion mutations (Methods). The first principal component separates steelhead trout from hatchery populations and cutthroat trout (Fig. 1D). The second principal component separates cutthroat trout and cutthroat hybrids from other trout. We performed an admixture analysis, with three clusters showing the lowest cross validation error (Figs. 1E, S2). I agreement with the PCA analysis, historical and contemporary steelhead cluster together, with cutthroat trout and hatchery trout forming the other two clusters.

All historical steelhead cluster near their corresponding contemporary population except for two groups. The first includes three historical South Fork Eel River (hereafter, Eel River) individuals that are genetically intermediate between steelhead and cutthroat trout. Considering both the PCA and ADMIXTURE analysis, there is strong genomic evidence that these historical Eel River fish are steelhead/cutthroat trout hybrids. The second historical group that is separate from its contemporary counterpart includes four San Lorenzo River individuals. Our ADMIXTURE analysis also indicates that these are either hatchery trout or steelhead-hatchery hybrids, which is consistent with historical records of an early hatchery at that location (see Discussion). Since these seven historical samples contain alleles from another species and from non-native domesticated trout, we removed them from further analyses to avoid introducing bias into our historical baseline.

Almost all trout in our five contemporary steelhead populations (Eel River, Coyote Creek, San Lorenzo River, Llagas Creek, and Nacimiento River) grouped with their historical populations (Fig. 1D), indicating that these contemporary steelhead populations derive most, if not all, of their ancestry from the historical steelhead populations. The only exceptions were four contemporary steelhead individuals, two from the Eel River and two from Nacimiento River, that appear to contain substantial hatchery trout ancestry based on both ADMIXTURE and PCA analysis.

Analysis of mitochondrial data was broadly consistent with nuclear data and previous research. We generated a tree of all samples using mitogenome data (Fig. S3) and found the tree structure generally reflected our nuclear genome analysis, but hatchery trout and coastal steelhead were less clearly distinguished with mtDNA, consistent with prior research^13^. Finally, we confirmed that the historical mitogenomes were consistent with the mtDNA haplotypes identified by Pearse et al. (Table S2).

### Measures of population structure show strong similarities between contemporary and historical steelhead

We found a strong pattern of isolation by distance in both historical and contemporary steelhead. We computed an outgroup f3 statistic (a measure of genetic similarity) for all pairs of historical populations and found it to be highly correlated with coastline distance between populations, confirming strong isolation by distance in the historical samples (black dots, Fig 2A; *ρ* = -0.98, p = 0.017). When we repeated this analysis using all contemporary samples, we again found a robust signal of isolation by distance (Fig 2B; *ρ* = -0.95, p = 0.008), consistent with a previous study of nuclear DNA markers encompassing most of the range of California steelhead that found strong contemporary isolation by distance^12^. These results are in contrast with prior work using a short mtDNA sequence in these same populations that found a strong pattern of isolation by distance in the historical collections only^13^. However, when we considered genetic distance between all pairs of individual fish (green and orange circles, Fig 2A,B), the variability is noticeably greater in the contemporary samples than in the historical samples, in large part due to the few hatchery hybrids present in contemporary samples (Fig 2B, orange circles).

**Fig. 2.**
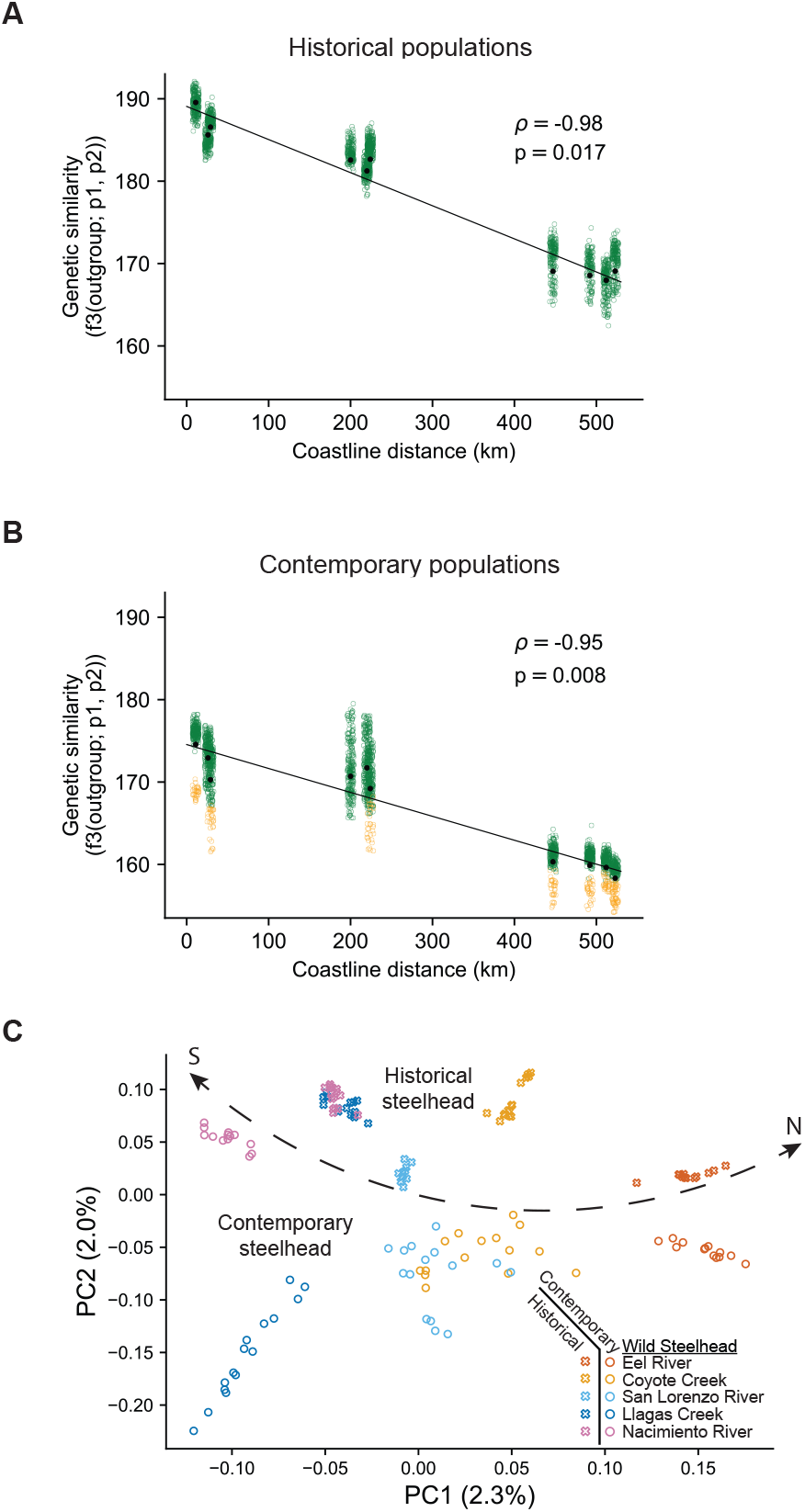
Isolation by distance and PCA suggest persistent North-South population structure. Isolation by distance calculated by outgroup f3 statistic between all pairs of populations (black dots) and all pairs of individual trout (green circles for non-hybrid pairs, orange circles for pairs that include at least one hybrid) in (**A**) historical populations and (**B**) contemporary populations. Best fit line, Pearson’s correlation, and Mantel test p-value are calculated using data from all pairs of populations (black dots). Jitter was added to the x-axis of the green circles to improve visualization. (**C**) A PCA of contemporary populations after removing trout with suspected hatchery or cutthroat trout ancestry. Dotted line separates historical and contemporary samples, with populations largely mirrored across the line.

When we removed all contemporary trout with suspected hatchery ancestry (two from Nacimiento River and two from Eel River) and performed a PCA analysis, contemporary population structure largely reflected historical population genetic and geographic structure (Fig. 2C). PC1 captured coastline distance, and PC2 roughly separated historical and contemporary populations.

### D-statistics reveal limited hatchery introgression into contemporary populations

We find a low but statistically significant amount of hatchery introgression in most contemporary populations. As described above, population structure analysis (Figs. 1D,E, 2C) suggests that many of the contemporary steelhead derive little ancestry from the hatchery strains we sampled. To explicitly test for introgression between hatchery strains and contemporary steelhead that occurred after historical steelhead were sampled, we calculated D-statistics of the form D(Chinook, hatchery, historical X, contemporary X,) where X is any of the five steelhead populations in this study, hatchery is any of the five hatchery strains we considered, and Chinook salmon is used as an outgroup. Positive D-statistics indicate excess allele sharing between a hatchery strain and a contemporary steelhead population, whereas negative D-statistics indicate excess allele sharing between a hatchery strain and a historical steelhead population (Fig. 3A). D-statistics revealed highly significant (|Z| > 6) introgression from hatchery strains into contemporary Eel River steelhead and into historical San Lorenzo River steelhead (Fig. 3B). In both of these populations, our PCA analysis found likely steelhead-hatchery hybrids, although those presumed hybrids were not included in this D-statistic analysis. We observed introgression from all hatchery strains into contemporary Eel River steelhead, and only from the Mt. Whitney, Mt. Shasta, and Coleman strains into the historical San Lorenzo River steelhead. These are the strains that are closest to the presumed historical San Lorenzo hatchery trout in the PCA (Fig. 1D). There was also a signal of moderately significant introgression (6 > |Z| > 3) into contemporary Llagas Creek, Nacimiento River, and San Lorenzo River steelhead. Since the D-statistic does not scale linearly with gene flow, it is difficult to quantify the fraction of hatchery gene flow into contemporary steelhead. For context, we note that these D-statistics are all lower than that found when calculating gene flow from neanderthals into non-Europeans.

**Fig. 3.**
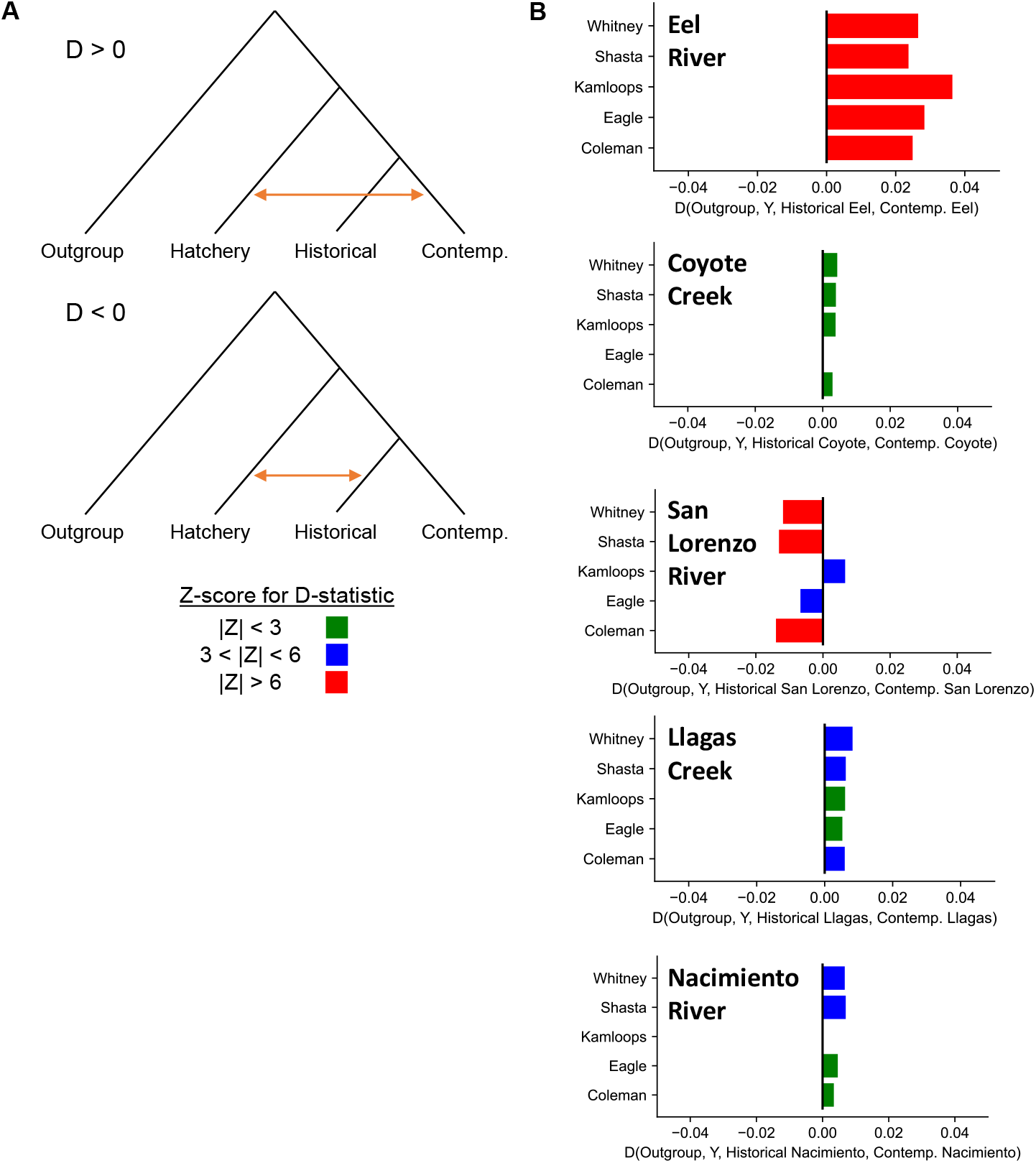
Contemporary steelhead show minimal hatchery introgression. (**A**) Diagram of D-statistic topology and interpretation, where a positive D-statistic indicates excess allele sharing between hatchery and contemporary populations, and a negative D-statistic indicates excess allele sharing between hatchery and historical populations. Z-scores greater than 3 are considered moderately significant, with Z-scores greater than 6 highly significant. (**B**) D-statistic for all five rivers, quantifying significance of introgression between historical or contemporary steelhead populations and any of the five hatchery strains. Z-scores colored according to (A).

### Contemporary steelhead show signs of selection on migration-associated locus

We found a consistent reduction over time in the frequency of a haplotype associated with migratory tendency in California steelhead^14^. We found that 90% of sampled historical trout were homozygous for the migration-associated haplotype, and only 1% were homozygous for the resident-associated haplotype (Fig. 4A). Since these historical trout were collected, all five rivers have been significantly modified by dams, water diversion, and habitat degradation, impeding steelhead migratory behavior. Overall, we found only 45% of contemporary trout were homozygous for the migration-associated haplotype, and 17% were homozygous for the resident-associated haplotype, even after removing samples with clear hatchery ancestry. All five rivers showed an increase in the resident-associated haplotype, with Llagas Creek having the greatest increase, and the San Lorenzo River having the least change. We note that these haplotypes are largely distinct in the hatchery rainbow trout strains (Fig. S4), which is consistent with previous work^14^ and indicates that this increase in resident-associated haplotypes is due to selection and not introgression.

**Fig 4.**
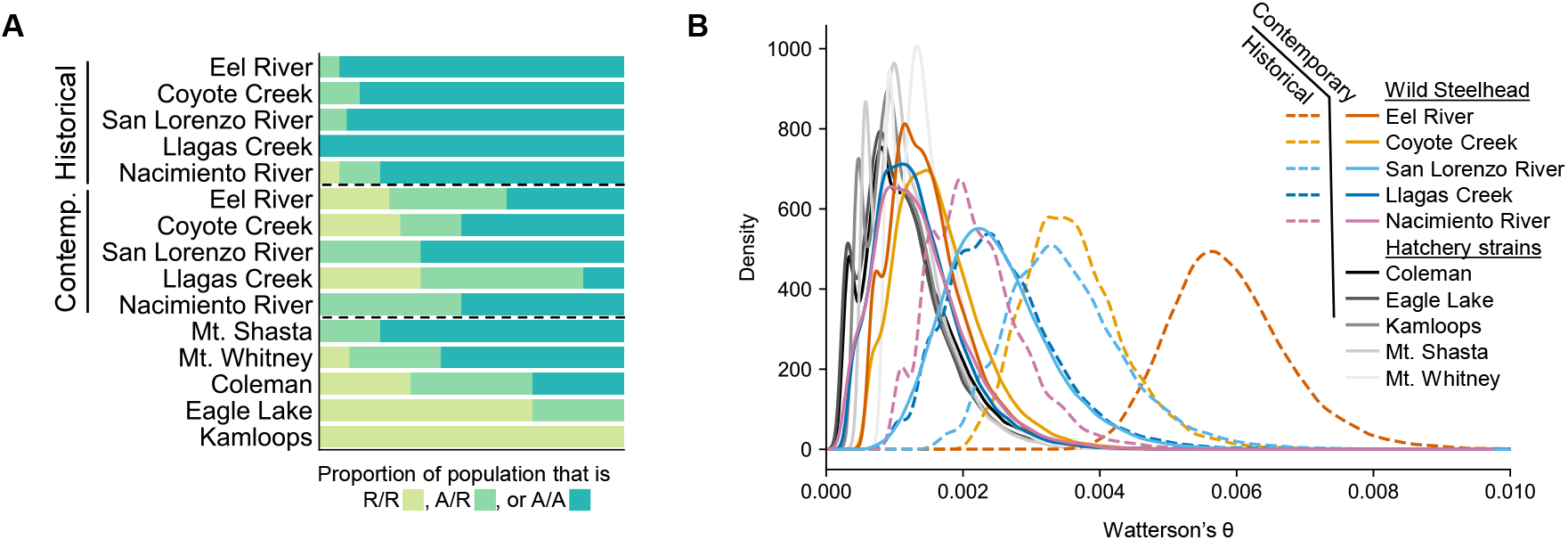
Contemporary steelhead genomes show signs of adaptation and lost genetic diversity. (**A**) Comparison of the proportion of the Omy05 migration-associated haplotype in historical and contemporary populations. R indicates the resident/rearranged haplotype and A indicates the anadromous/ancestral haplotype. (**B**) Comparison of genomic diversity present in historical (dotted lines), contemporary (solid lines), and hatchery samples (solid gray lines). Contemporary and historical hatchery hybrids were not included in the analysis.

### Genomic diversity has decreased over time

Comparison of Waterson’s theta and pi across historical and contemporary steelhead populations revealed a clear signal of reduced genomic diversity over time (Figs. 4B, S5). Among historical populations, the Eel River had the highest genomic diversity, which is presumably a consequence of higher long-term effective population size or cutthroat trout introgression. Additionally, we found that in our historical populations, northern populations showed greater genetic diversity than southern populations. Among contemporary populations, the San Lorenzo River had the highest genomic diversity, on par with the historical genomic diversity seen in the two southernmost historical populations. The remaining contemporary populations had similar levels of genetic diversity, in contrast to the pattern of increasing genetic diversity with latitude observed in historical populations.

## Discussion

Given the immense environmental pressures that California steelhead trout have faced in the past century, there is reasonable concern that their population genomic structure and ancestry has been radically altered. Although the effects of stocking with domesticated hatchery trout varied by river, we found that contemporary steelhead populations were all genetically similar to their historical counterparts. Introgression analysis found statistically significant, although relatively minimal, hatchery introgression in contemporary Eel River, Llagas Creek, and Nacimiento River populations. Finally, we found a dramatic decrease over time in the frequency of a haplotype associated with migration.

Despite widespread habitat loss, dam construction, and hatchery stocking throughout the 20th century, we find that steelhead population structure has changed little in the past century. We found significant isolation by distance in both historical and contemporary steelhead populations. This result supports the conclusions of Garza et al. (2014) that isolation by distance persists in contemporary steelhead populations. These results differ from those found by Pearse et al. (2011), and this is likely explained by our use of whole genome data, which is able to distinguish diversity that is missed in a mitochondrial haplotype analysis. Given this isolation by distance, along with concordant results from our ADMIXTURE (Fig. 1C) and PCA analysis (Fig. 2C), we conclude that the present steelhead population structure strongly resembles the structure from the turn of the twentieth century.

Researchers have long debated the extent to which contemporary steelhead populations may be introgressed with domesticated hatchery rainbow trout and how this may be affected by the presence of barriers to migration. Our results suggest that the effects of hatchery stocking were largely similar across rivers, with no or low or levels of hatchery hybridization and introgression. However, the effects of hatchery stocking were subtly different between rivers in unpredictable ways. For example, our contemporary samples from the South Fork Eel River, often considered relatively pristine steelhead habitat, had the strongest signal of hatchery introgression and included two clear steelhead-hatchery trout hybrids. While this is unexpected, it is plausible, since 9–17 million trout were planted in the Eel River over the last century, with the South Fork Eel River receiving large numbers of hatchery trout through 1995^22^. In the comparably urban Coyote Creek, which passes through San Jose, a major city with over one million residents, we found no detectable introgression, despite heavy hatchery stocking of Coyote Reservoir, upstream of the collection site.

The surprisingly low amount of hatchery introgression observed may be explained by evolutionary selection. Hatchery trout are artificially selected to thrive in a hatchery environment, produce eggs throughout the year^3^, and generally lack genetic variation^6,11^, including alleles associated with local adaptation in native steelhead and rainbow trout^23^. These factors may lead to poor survival or mating success for hatchery trout released into the wild^23^. For example, we observed hatchery introgression into historical San Lorenzo steelhead, but this introgression was lessened in contemporary San Lorenzo steelhead. This disappearance of hatchery introgression suggests that pulses of hatchery admixture may eventually be mitigated, possibly through selection against alleles that were selected for in hatchery breeding but confer low fitness in the wild. Soon after our contemporary samples were collected, the California Department of Fish and Wildlife transitioned towards stocking only sterile (triploid) fish in reservoirs connected to native trout populations, and thus further hatchery influence is unlikely, although not impossible.

Among the historical samples, we observed two types of hybrids that are consistent with the historical record. The first type were hatchery hybrids in our historical San Lorenzo steelhead. These are consistent with historical records of an operational fish hatchery that released trout into the San Lorenzo River beginning in 1905, and possibly earlier. The hatchery hybrids and introgression we observed suggest that the hatchery was relatively successful, contributing to the steelhead/rainbow breeding population. Indeed, when the historical samples were collected, the researchers noted that a few specimens were distinct, with larger eyes and a different color pattern^24^. Despite a century passing, contemporary hatchery strains appear genetically similar to these historical hatchery strains, particularly Mt. Whitney and Coleman (Fig. 1D).

The second type of hybrids that align with the historical record were cutthroat trout-steelhead hybrids in historical samples from the Eel River. These are anticipated, as the Eel River marks the southern extent of the geographic range of cutthroat trout. Indeed, we found a signal of residual cutthroat trout ancestry in all Eel River steelhead individuals, which is consistent with other work that has found that the two species hybridize wherever they are sympatric, but that introgression is limited and does not typically result in a large fraction of fit hybrids (i.e. a hybrid swarm)^25,26^.

We found a striking reduction in the frequency of the Omy05 migration-associated haplotype across all steelhead populations, even those below barriers. It had previously been shown that above-barrier populations had a greater proportion of resident-associated alleles compared to their below-barrier counterparts^15,27^. This increase is attributed to a rapid adaptation to the migratory constraints of trout trapped above barriers. Our results show that not only above-barrier, but also below-barrier populations have shifted significantly from their pre-disturbance baseline, in which 90% of historical fish sampled were homozygous for the migration-associated haplotype. For example, both the Eel River and Coyote Creek were sampled below a barrier and showed a 30-40% increase in the frequency of the resident-associated allele over time. Given that this shift was previously unknown and management plans have been based on contemporary allele frequencies, we contend that this is a rare validated example of Pauly’s shifting baseline syndrome for genomic traits in fish. This change from the historical baseline suggests that steelhead are evolving towards a more frequent resident life history even when migration is not blocked by complete barriers, possibly in response to habitat loss, reduced river discharge, or partial barriers ^28^. Alternatively, after above-barrier populations experience purifying selection favoring resident haplotypes, some of these fish may pass downstream through barriers. They will not be able to return above the barrier to reproduce, and may be contributing to the higher frequency of resident-associated haplotypes^29^.

While it’s clear that this shift in migration-associated haplotypes is occurring, the degree of change varies greatly across rivers. The two rivers with the least change in migration-associated haplotype over time were the San Lorenzo River and the Nacimiento River. The San Lorenzo is our only site not directly affected by a dam during its history, and it also shows the highest nucleotide diversity of our contemporary samples. In contrast, the Nacimiento River has been heavily impacted by a dam. One possible explanation for the high frequency of migration-associated alleles may be that the Nacimiento reservoir is so large that fish are able to complete an adfluvial migration, migrating between the reservoir and upstream tributaries without ever reaching the ocean. Indeed, prior work has found that the migration-associated haplotype is at higher frequency in above-barrier populations in river systems with larger reservoirs^27^. Research to better understand why the haplotype frequency varies widely between contemporary populations may help inform management plans. These plans should incorporate the discovery that the migration-associated haplotype has decreased in frequency over time and emphasize actions that will increase opportunities for anadromous migration.

Our results indicate that conservation efforts to protect present-day populations can effectively preserve historical population structure. Compared to their historical abundance, most steelhead populations in California, Oregon, and Washington have declined substantially. Starting in the 1800s, logging and hydraulic mining degraded or destroyed large regions of steelhead habitat.

Given these early impacts, it is likely that our historical samples do not represent a completely pre-disturbance baseline, but they do provide a glimpse of this species prior to most modern changes. Throughout the 1900s, habitat availability and quality continued to decrease due to dam construction, water diversion, pollution, and a dramatic expansion of hatchery stocking. As a consequence, many steelhead are listed as threatened or endangered under the US Endangered Species Act (ESA). Since hatchery stocking has had relatively minimal impact on population structure, effective conservation strategies to increase population sizes, such as restoring habitat^30^, should continue to be prioritized.

Although the steelhead populations we studied here are protected under the ESA, their conspecifics above impassable barriers are not protected, since they are considered to be rainbow trout. Our findings of low levels of hatchery introgression in the above-barrier Llagas Creek and Nacimiento River population contributes to growing evidence that many of the fish of coastal steelhead lineage trapped above barriers are genetically very similar to the anadromous steelhead downstream^11,27^. We emphasize that effective conservation strategies need to recognize that fish that travel down a barrier may then contribute genetically to below-barrier populations ^15,29,31^(Kobayashi et al. in press).

In this work, we investigate a limited number of rivers and fish per river, as our study design was constrained by the scope of the historical collections. Even with these modest sample sizes, we would have been able to observe large genomic changes present in these populations. Additionally, these rivers were stocked with several hatchery strains over the past century. The strains that we sampled are the dominant ones in use in California over the last half century and are representative of major domestication lines^3^. While we cannot rule out that other strains could have contributed to introgression, we expect that the sampled strains are representative of possible introgression sources.

Our results underscore the pivotal role that museum collections continue to play in our understanding of how populations may have shifted from their pre-disturbance baseline. Such collections provide a unique opportunity to calibrate the current state of a species through direct comparison with historical populations. Even samples that are often considered poor candidates for historical DNA sequencing, such as those stored in formalin, may yield valuable genetic information as techniques continue to improve. The consistency of our results with historical records demonstrates this and indicates that we took appropriate steps to limit known biases from historical DNA, such as those arising from cytosine deamination and genome mapping methods. For species without available museum collections, remains preserved in indigenous middens or other sub-fossil deposits may provide an alternative source of historical DNA. Although historical collections of populations (i.e., many conspecifics sampled simultaneously), such as those used in this study, are rare, there remain a large number of individual historical specimens that we hope will be leveraged to improve conservation practices today.

## Methods

### Samples, library prep, and sequencing

Historical samples were obtained as described in ^13^. Briefly, we obtained fin clip tissue samples of *O. mykiss* (identified as *Salmo irideus* or *S. gairdneri)* from 75 specimens^24^ stored at the Smithsonian Institution’s National Museum of Natural History that were collected from five rivers in 1897 and 1909 (Supplementary table 1, Fig. 1A-C). These specimens are currently stored in ethanol, although they were almost certainly initially preserved in formalin. Historical sample DNA extraction was done in an isolated laboratory facility, occupied by the authors since construction. Equipment was cleaned with bleach prior to use to avoid contamination with contemporary DNA. We extracted DNA from each sample using the DNeasy 96 tissue protocol (Qiagen, Inc.), with two negative extraction controls included in each 96-well extraction plate. Prior to library preparation, we quantified extracts using the Qubit dsDNA HS kit (Invitrogen) and a Qubit 4 fluorometer (Invitrogen), we estimated the ssDNA molar concentration using an estimated fragment length of 60bp.

For historical samples, single stranded libraries were prepared from extracted DNA according to ^20^ with the following modifications for Llagas Creek: a final reaction volume of 80 μl; a 2:1 ratio of splinter:adaptor oligonucleotide; a 6:1 molar ratio of adaptor:ssDNA; and a 65:1 molar ratio of extreme thermostable single-stranded DNA binding protein:ssDNA. We double-indexed and amplified these libraries and prepared 50 μl reactions containing 10 μl pre-amplified library, 1X AmpliTaq Gold 360 Master Mix (Applied Biosystems), 1 μM i7 indexing primer and 1 μM i5 indexing primer. We amplified libraries at 95° C for 10 m, followed by 20 cycles of 95° C for 30 s, 60° C for 30 s and 72° C for 60 s, followed by 72° C for 7 m. For samples from Nacimiento River, South Fork Eel River, Coyote Creek and San Lorenzo River, we made modifications as described in the supplementary materials of ^32^. Extraction and library controls were processed alongside samples in order to detect possible contamination. All libraries received unique non-overlapping dual indices to reduce the likelihood of index hopping. All historical samples were sequenced on a NovaSeq 6000 or HiSeq X.

Contemporary samples were collected from the same five rivers between 1997 and 2004 (Supplementary Table 1, Fig. 1) and are described by ^11,15^. We extracted genomic DNA from caudal fin clips that had been air-dried on blotter paper and stored at room temperature using the DNeasy 96 Blood and Tissue Kit on a BioRobot 3000 (Qiagen Inc.). Extractions were selected for whole-genome sequencing prep based on both sample quality, assessed via agarose gel electrophoresis, and concentration, quantified using the Qubit Flex Fluorometer using the dsDNA Broad Range Assay Kit (Thermo Fisher Scientific). For all samples from hatchery strains (Coleman, Eagle Lake, Kamloops, Mt. Shasta, and Mt. Whitney) as well as contemporary samples from Coyote Creek, South Fork Eel River and Llagas Creek, libraries were prepared from extracted DNA using a library preparation previously described by ^33^. These libraries were sequenced on a HiSeq X. For the remaining contemporary samples (Nacimiento River and San Lorenzo River), libraries were prepared using the NEBNext Ultra II FS DNA Library Prep Kit for Illumina to prepare libraries, along with NEBNext Unique Dual Index primer pairs to barcode each sample. Validation of library construction success was done using the Qubit dsDNA HS Assay kit (Thermo Fisher Scientific) to measure concentration and the TapeStation with the High Sensitivity D10000 ScreenTape assay (Agilent) to confirm quality. Libraries were then normalized, pooled, and sent for sequencing at the North Carolina State University Genomic Sciences Laboratory on an Illumina NovaSeq 6000 using an S4 flow cell and paired-end 150bp chemistry targeting 2X coverage for each sample.

### Data Analysis

Raw fastq files were processed with SeqPrep2 (https://github.com/jeizenga/SeqPrep2) to trim adapters and merge reads, with the following arguments: -q 15 -L 30 -o 15 -m 0.05 -A AGATCGGAAGAGCACACGTC -B AGATCGGAAGAGCGTCGTGT -C ATCTCGTATGCCGTCTTCTGCTTG -D GATCTCGGTGGTCGCCGTATCATT -S. For historical samples, bwa aln ^34^ was used to align both merged and unmerged reads to the OmykA_1.1 genome ^35^, using the following arguments: -l 16500 -n 0.01 -o 2. For contemporary samples, bwa mem ^36^ with -M option was used to align both merged and unmerged reads to the OmykA_1.1 genome. Aligned merged and unmerged reads were combined into a single bam file using samtools merge. Picard ^37^ MarkDuplicates v2.21.7 was used to remove duplicates from the combined bam with the following arguments: REMOVE_DUPLICATES=true MAX_FILE_HANDLES_FOR_READ_ENDS_MAP=800 VALIDATION_STRINGENCY=LENIENT.

Single nucleotide variants with a minor allele frequency of 5% or greater in the superpopulation consisting of all contemporary, historical, and hatchery *O. mykiss* samples were then identified using angsd v0.935-52-g39eada3 ^38^. The following options were used: -uniqueOnly 1 -remove_bads 1 -trim 0 -C 50 -minMapQ 20 -minQ 20 -minInd 50 -setMinDepth 50 -setMaxDepth 2000 -doCounts 1 -GL 2 -doGlf 2 -doMajorMinor 4 -doGeno 3 -doPost 2 -doMaf 1 -minMaf 0.05 -nInd 200 -rmTrans 1 -only_proper_pairs 0. We removed transitions due to their high error rate in historical DNA ^39^.

Since the *O. mykiss* genome has high repeat content, we removed regions of the genome that had poor mappability, in order to reduce artifacts in downstream analysis. We first used GenMap v1.2.0 ^40^ to compute the (30,2)-mappability (uniqueness of 30 bp-mers with up to two mismatches) in the OmykA_1.1 genome. We then identified every position in the genome for which all possible overlapping 40 bp (or shorter) reads have a (30,2)-mappability of 1 (i.e., are unique). Next, we ran RepeatMasker ^41^ using a custom repeat library ^35^ and ‘slow search’ (-s) option. After filtering, we had 981,618 single nucleotide variants remaining that were used in all subsequent analysis.

To reduce reference bias, we then returned to our mapped reads for each of the historical samples and performed additional filtering using scripts developed by Günther and Nettelblad (2019). We used the modify_read_alternative.py script to first swap the reference and alternate allele in all mapped reads that overlap a single nucleotide variant. These altered reads are then re-mapped to the OmykA_1.1 genome using the bwa aln parameters described above, using a modified version of the remap_modified_reads.sh script. We then used the filter_sam_startpos_dict.py script to identify reads that no longer map to the same location in the genome, and remove from the alignment the equivalent unswapped read.

We used pileupCaller ^43^ to call pseudo-haplotypes for each sample using all reads that passed filtering. Omy05 was removed to avoid bias due to the change in frequency of migration-associated alleles over time. Calling pseudo-haplotypes (selecting a random read at all variant sites) is commonly used when historical DNA coverage is too low to confidently call genotypes ^44^. Plink v1.90b6.26 ^45^ was then used to prune variants in linkage equilibrium using the following parameters: --indep-pairwise 50 5 0.5. SmartPCA v18140^46^ was used to generate the PCA, with the following parameter: outliersigmathresh:12. Admixtools^47^ qpDstat v980 was used to generate D-statistics. Admixtools qp3pop v651 was used to generate outgroup f3 statistics with parameter outgroupmode: YES. ADMIXTURE v1.3.0 ^48^ was used to generate ADMIXTURE plots using the –cv parameter for K values from 1 to 5.

We downloaded Chinook salmon sequencing reads from SRA samples SRR12798421, SRR12798422, SRR12798424, SRR12798425, SRR12798426, SRR12798427, SRR12798349, SRR12798351, and SRR12798352 ^49^. We downloaded cutthroat sequencing reads from SRA samples SRR16192176–SRR16192181 from PRJNA402066. These reads were processed, aligned, and called in the same way as contemporary *O. mykiss* samples, described above.

To create the mitochondrial tree, we used processed reads (as described above). Mia (https://github.com/mpieva/mapping-iterativeassembler) was then used to determine a consensus mitochondrial sequence for each sample, using the Arlee reference mitochondrion (RefSeq: NC_001717.1). To avoid calling bases that could be the result of ancient DNA damage, we required a minimum of three agreeing reads to call a base at each site, and 2/3 agreement between mapped reads that exceeded the minimum 3x coverage. Sites not meeting these criteria were classified as N. MUSCLE v5.1 ^50^ was used to align all consensus mitochondrial sequences using default parameters. RAxML v8.2.12 ^51^ was used to run a rapid bootstrap analysis and search for the best-scoring ML tree with a GTRGAMMAI model and 1000 bootstop permutations (full command: raxml -f a -x 43 -p 42 -# autoMRE -m GTRGAMMAI --bootstop-perms 1000 -s input.afa -n output.raxml). Toytree ^52^ was used to visualize the tree.

To generate isolation by distance figures, we calculated coastline distance between pairs of rivers by considering potential marine routes steelhead could have taken between rivers. Specifically, these routes began at the point where a river drains to the ocean and were constructed using a series of straight lines that remained within 20 miles of the coast. We used the stats.linregress function from SciPy v1.7.3 ^53^ to calculate best fit line, and used the test function from the python package mantel v2.2.1 (https://github.com/jwcarr/mantel) to perform a Mantel test to calculate Pearson correlation and p-value.

To calculate theta and pi for each population, we followed Korneliussen et al.^54^. Briefly, we ran ANGSD on each population with the following options: -uniqueOnly 1 -remove_bads 1 -only_proper_pairs 1 -trim 5 -C 50 -minMapQ 20 -minQ 20 -setMaxDepth 90 -noTrans 1 -doCounts 1 -GL 2 -doSaf 1. We also limited analysis to all sites that passed the GenMap mappability analysis described above. Next, we used ANGSD realSFS to generate a folded SFS for each population, followed by saf2theta to calculate theta/pi for each site. Then we ran ANGSD do_stat to calculate theta/pi across a sliding window of 50kb and a step of 10kb. For each window, we then calculated a normalized theta/pi by dividing theta/pi by the number of sites in the window with data. Finally, we plotted the kernel density estimation for normalized theta/pi over all windows that had more than 500 sites with data.

## Acknowledgements

We thank Beth Nelson, Elena Correa, Edith Martinez, Sarah Ford, Ezra Collins, Delight Lee, Halle Bender, Josh Kapp, Jeff Rodzen, and Jonas Oppenheimer for their contributions to this project. We thank all members of UCSC Paleogenomics Lab for their insightful feedback. AS was supported by an NSF PRFB Program under Grant No. 2109912.

