## Supplementary Figures for "Recent adaptation in an imperiled salmonid revealed by museum genomics"

### Supplementary Material for “Recent adaptation in an imperiled trout revealed by museum genomics”

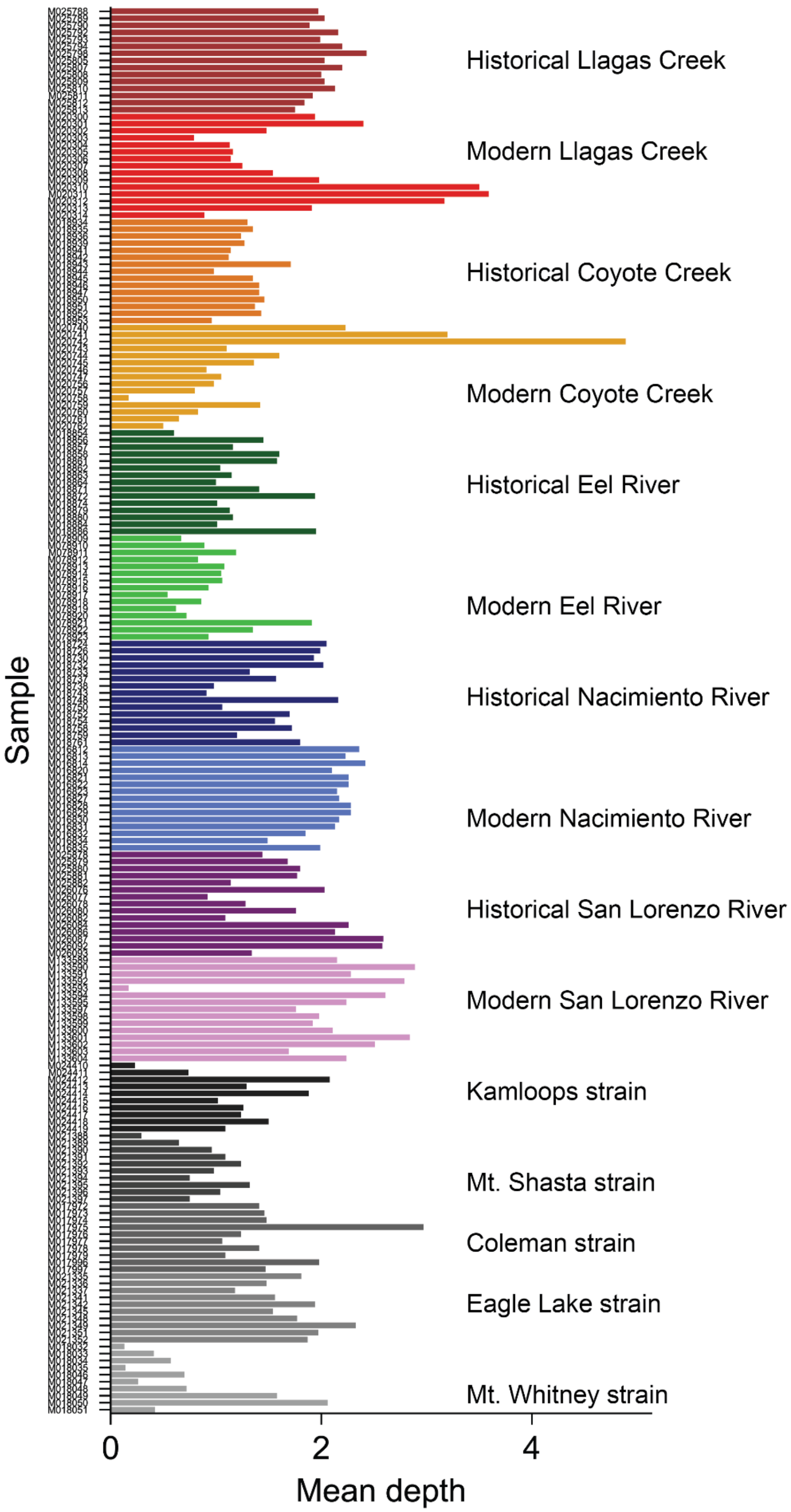
Historical Llagas Creek
Modern Llagas Creek
Historical Coyote Creek
Modern Coyote Creek
Historical Eel River
Modern Eel River
Historical Nacimiento River
Modern Nacimiento River
Historical San Lorenzo River
Modern San Lorenzo River
Kamloops strain
Mt. Shasta strain
Coleman strain
Eagle Lake strain
Mt. Whitney strain
Sample
0
2
4
Mean depth

Fig. S1. Depth of coverage for all 200 samples sequenced in this study. Sample names correspond to the “NMFS_DNA_ID” column in Table S1.

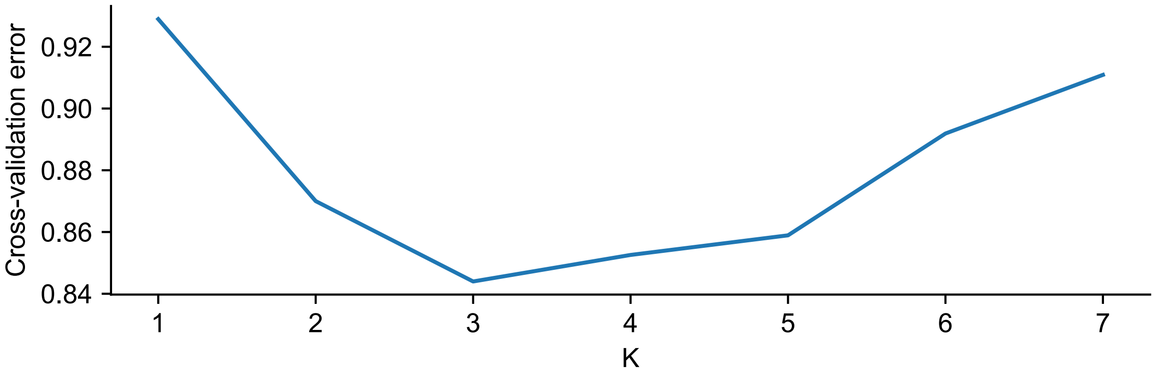


Fig. S2. Cross-validation error as we vary the number of clusters (K) in our ADMIXTURE analysis.

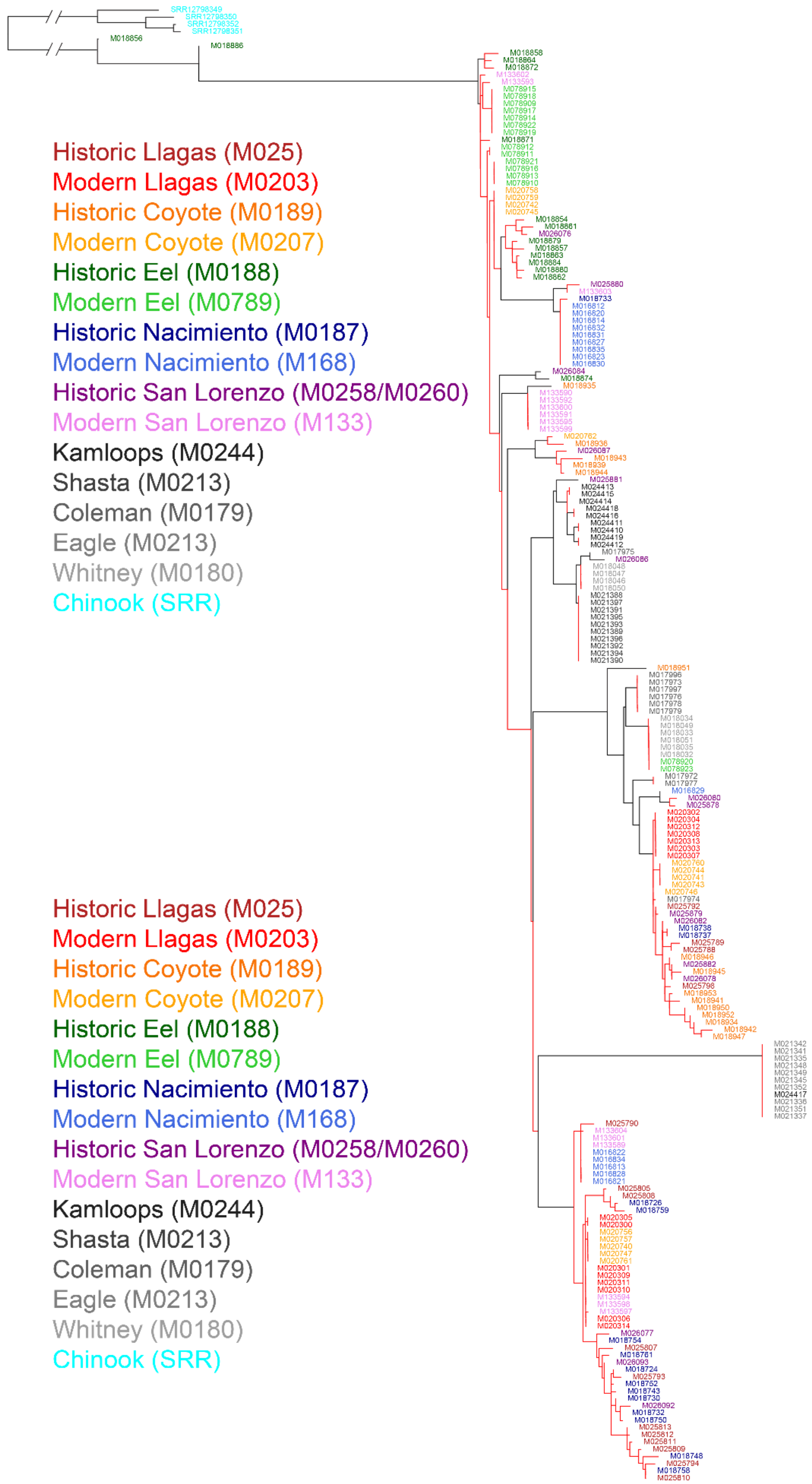
Historic Llagas (M025)
Modern Llagas (M0203)
Historic Coyote (M0189)
Modern Coyote (M0207)
Historic Eel (M0188)
Modern Eel (M0789)
Historic Nacimiento (M0187)
Modern Nacimiento (M168)
Historic San Lorenzo (M0258/M0260)
Modern San Lorenzo (M133)
Kamloops (M0244)
Shasta (M0213)
Coleman (M0179)
Eagle (M0213)
Whitney (M0180)
Chinook (SRR)
Historic Llagas (M025)
Modern Llagas (M0203)
Historic Coyote (M0189)
Modern Coyote (M0207)
Historic Eel (M0188)
Modern Eel (M0789)
Historic Nacimiento (M0187)
Modern Nacimiento (M168)
Historic San Lorenzo (M0258/M0260)
Modern San Lorenzo (M133)
Kamloops (M0244)
Shasta (M0213)
Coleman (M0179)
Eagle (M0213)
Whitney (M0180)
Chinook (SRR)

Fig. S3 Maximum likelihood phylogenetic tree of mitochondrial genomes. The tree was rooted using Chinook salmon mitochondria. Red lines indicate <90% bootstrap support. Codes given after names are the first 4-5 characters of the IDs in the tree for that group, in order to aid matching when colors are ambiguous.

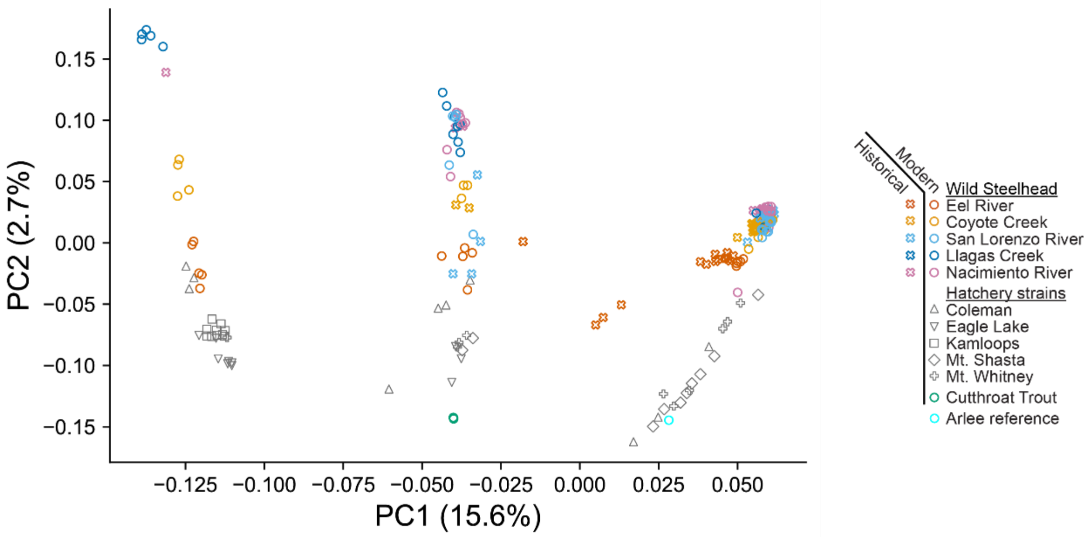


Fig. S4. PCA generated by considering only those variants on omy05 between positions 32,000,000 and 85,000,000, capturing a double inversion that is associated with migration tendency. The three clusters on PC1 correspond to homozygous resident (R/R) on the left, heterozygous (A/R) in the center, and homozygous anadromous (A/A) on the right. PC2 largely separates steelhead (above) from hatchery trout (below).

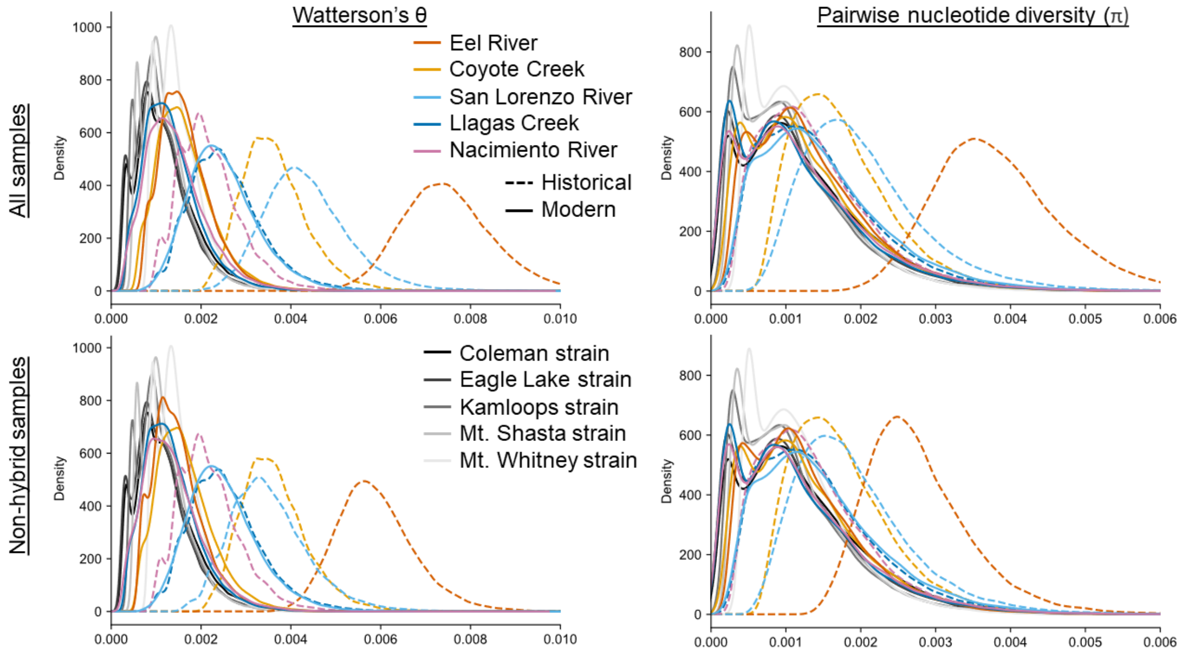


Fig. S5. Distribution of genomic diversity in modern, historical, and hatchery samples. Hatchery samples are shown in shades of gray. The bottom plots are calculated using only samples that show no clear signs of hybridization with cutthroat or hatchery trout in our PCA analysis. The bottom left plot is identical to Fig. 4B.
